# Multicellular Spatial Model of RNA Virus Replication and Interferon Responses Reveals Factors Controlling Plaque Growth Dynamics

**DOI:** 10.1101/2021.03.16.435618

**Authors:** Josua O. Aponte-Serrano, Jordan J.A. Weaver, T.J. Sego, James A. Glazier, Jason E. Shoemaker

## Abstract

Respiratory viruses present major health challenges, as evidenced by the 2009 influenza pandemic and the ongoing severe acute respiratory syndrome coronavirus 2 (SARS-CoV-2) pandemic. Severe RNA virus respiratory infections often correlate with high viral load and excessive inflammation. Understanding the dynamics of the innate immune response and its manifestation at the cell and tissue levels are vital to understanding the mechanisms of immunopathology and developing improved, strain independent treatments. Here, we present a novel spatialized multicellular spatial computational model of two principal components of tissue infection and response: RNA virus replication and type-I interferon mediated antiviral response to infection within lung epithelial cells. The model is parameterized using data from influenza virus infected cell cultures and, consistent with experimental observations, exhibits either linear radial growth of viral plaques or arrested plaque growth depending on the local concentration of type I interferons. Modulating the phosphorylation of STAT or altering the ratio of the diffusion constants of interferon and virus in the cell culture could lead to plaque growth arrest. The dependence of arrest on diffusion constants highlights the importance of developing validated spatial models of cytokine signaling and the need for *in vitro* experiments to measure these diffusion constants. Sensitivity analyses were performed under conditions creating both continuous plaque growth and arrested plaque growth. Findings suggest that plaque growth and cytokine assay measurements should be collected during arrested plaque growth, as the model parameters are significantly more sensitive and more likely to be identifiable. The model’s metrics replicate experimental immunostaining imaging and titer based sampling assays. The model is easy to extend to include SARS-CoV-2-specific mechanisms as they are discovered or to include as a component linking epithelial cell signaling to systemic immune models.

**Author Summary:** COVID-19 is possibly the defining healthcare crisis of the current generation, with tens of millions of global cases and more than a million reported deaths. Respiratory lung infections form lesions in the lungs, whose number and size correlate with severity of illness. In some severe cases, the disease triggers a severe inflammatory condition known as cytokine storm. Given the complexity of the immune system, computational modeling is needed to link molecular signaling at the site of inflection to the signaling impact on the overall immune system, ultimately revealing how severe inflammatory conditions may emerge. Here, we created a computational model of the early stages of infection that simulates lung cells infected with RNA viruses, such those responsible for COVID-19 and influenza, to help explore how the disease forms viral plaques, an *in vitro* analog to lesion growth in the lung. Our model recapitulates *in vitro* observations that pretreatment of biological signaling molecules called with type-I interferons, which are currently being evaluated for treatment of COVID-19. Analyzing the model, we, can stop viral plaque growth. We found that enhancing certain aspects of the innate immune system, such as the JAK/STAT pathway, may be able to stop viral plaque growth, suggesting molecules involved in this pathway as possible drug candidates. Quantifying the parameters needed to model interferon signaling and viral replication, experiments should be performed under conditions that inhibit viral growth, such as pretreating cells with interferon. We present a computational framework that is essential to constructing larger models of respiratory infection induced immune responses, can be used to evaluate drugs and other medical interventions quickly, cheaply, and without the need for animal testing during the initial phase, and that defines experiments needed to improve our fundamental understanding of the mechanisms regulating the immune response.

## Introduction

Respiratory virus infections contribute significantly to death rates worldwide. Seasonal influenza virus infection is responsible for 290,000 – 650,000 average annual deaths globally (1), and occasional, highly pathogenic pandemics strains can emerge, such as the 1918 Spanish Flu (2), resulting in significantly higher mortality rates. As of February 16^th^, 2021,November 2020, the SARS-CoV-2 virus has caused over 108.3 million recorded infections and 2.3 million deaths worldwide (3). Both influenza and SARS-CoV-2 are RNA viruses, and studies of severe SARS-CoV-2 and influenza infections find that impaired innate immune responses correlate with more severe outcomes (4–6). In highly pathogenic infections, aberrant innate immune responses – specifically elevated inflammation and high production of type-I interferons leading to hypercytokinemia (cytokine storm) (7) – are believed to be significant drivers of mortality (8,9). Excessive inflammation exacerbates tissue damage and hinders clinical recovery (10,11). Influenza studies show that immunomodulation can improve infection outcomes. Prestimulation of toll-like receptors is protective against highly pathogenic influenza strains in mice (12), while cell culture prestimulation with type-I interferons prevents viral plaque growth by SARS-CoV (13), SARS-CoV-2 (14), and influenza (15). Nebulized interferon α2b and interferon β are being investigated as an early treatment for COVID-19 (16,17). Collectively, these studies demonstrate the necessity of immune response regulation as a means to balance tissue damage from inflammatory responses with viral clearance. However, we cannot currently predict how an individual patient’s immune system will respond to a particular dosage and timing of an immunomodulator. Computational modeling may reveal how complex responses emerge during infection, rapidly identify and optimize treatments, and reveal dynamic insights into immunoregulation during respiratory infection that can drive the discovery and design of immune targeted treatments.

Recent computational models have considered many aspects of virus infection induced immune responses (18–21), however, few models describe virus replication and interferon regulation in a cell culture. Ordinary Differential Equation (ODE) based models assume either homogeneity or a compartment based structure, and typically ignore the diffusion of virions and cytokine signaling, heterogeneity of cell response to stimuli, and stochasticity of individual cells’ response (19,22). Many recent models (20,23,24) of interferon response use a simple virally resistant cell state which does not capture interferon stimulated genes’ (ISGs) effect on viral growth (23,24). A spatial model of viral spread and plaque growth (22) mentions the impact of diffusion constants on viral plaque formation, but does not incorporate the cells’ innate immune response to infection and paracrine induced activation, limiting the ability of the model to explain plaque growth arrest due to ISGs.

We developed a multiscale, multicellular, spatiotemporal model of the innate immune response to RNA viral respiratory infections *in vitro* in CompuCell3D (25) (CC3D), with the ability to simulate plaque growth, cytokine response, and plaque arrest. The model of IFN production and virus replication determines the conditions leading to arrest or promotion of plaque growth during the infection of lung epithelial cells with an RNA virus. Plaque growth assays are commonly used to compare virus growth rates across cell lines (22,26), quantify the number of infective agents (27,28), and observe the effects of drugs and compounds on virus spread (29–32). Plaque growth assays seed the virus at low multiplicity of infection (MOI) and allow it to replicate across a sheet of host cells in a cell culture to form visible plaques. Our aim in replicating plaque growth experiments in a spatial computer simulation was twofold. First, generating realistic simulations of physical experiments allows *in silico* experimentation for cheaper, faster, higher throughput hypothesis testing of more simultaneous outputs than would be experimentally viable (*i*.*e*. recording all species, continuous plaque growth and cell state information without staining or photographing). Second, our model replicates familiar biological experimental measurements and imaging, making our results accessible to wet lab biologists.

Our model includes two competing mechanisms, viral replication and the host cells’ innate immune response. Viral reproduction consists of the virus production within and export from infected cells, and viral particle spread via diffusion in the extracellular matrix. The host immune response includes interferon production, export, and diffusion, and the initiation of virally resistant cell states via ISGs. The model represents a monolayer of immobile human bronchial epithelial cells (HBECs). Each cell contains a separate model of epithelial immune signaling, viral replication and export, and cell death, based on a set of previously calibrated ODEs (33), modified to include species export to the extracellular environment. We adapt an existing model of cell transitions during viral infection (34) between susceptible, eclipse phase, virus releasing, and dead cell states. The extracellular environment allows diffusion of virions to spread the virus and form viral plaques and type-I interferons responsible for spatiotemporal paracrine signaling. The model gives insight into which processes are most important to IFN regulation and arresting the growth of viral plaques *in vitro* via IFN mediated antiviral cell states. Since epithelial regulation of IFN is an important early regulator of the immune system more broadly, the model could be expanded in future work to account for additional regulation via immune cells.

## Materials and Methods

The multicellular spatial model incorporates an ODE representation of epithelial cell cytokine production in response to RNA virus infection (35) into a field of agents using CompuCell3D (CC3D). This model simulates the replication and spread of virus within a monolayer of epithelial cells, and the cells’ attempts to arrest this viral spread through interferon signaling and ISG induced antiviral states within the cells.

### Cell States and Rationale

Fig. 1 provides a conceptual overview of the model. The model contains cells (agents) in 4 distinct states: uninfected (U), eclipse phase (I1) in which cells can produce IFN and virus replication occurs but there is no viral export, virus releasing (I2), wherein IFN is produced and virions are produced and exported, and dead (D), in which cells do not interact with their surroundings. The transition between each state is stochastic and depends on the cell’s local extracellular field concentration of virus and the health (H; described in Intracellular Model Equations and Rationale) of the cell, and defined by the following equations, adapted from (34):

**Fig 1.**
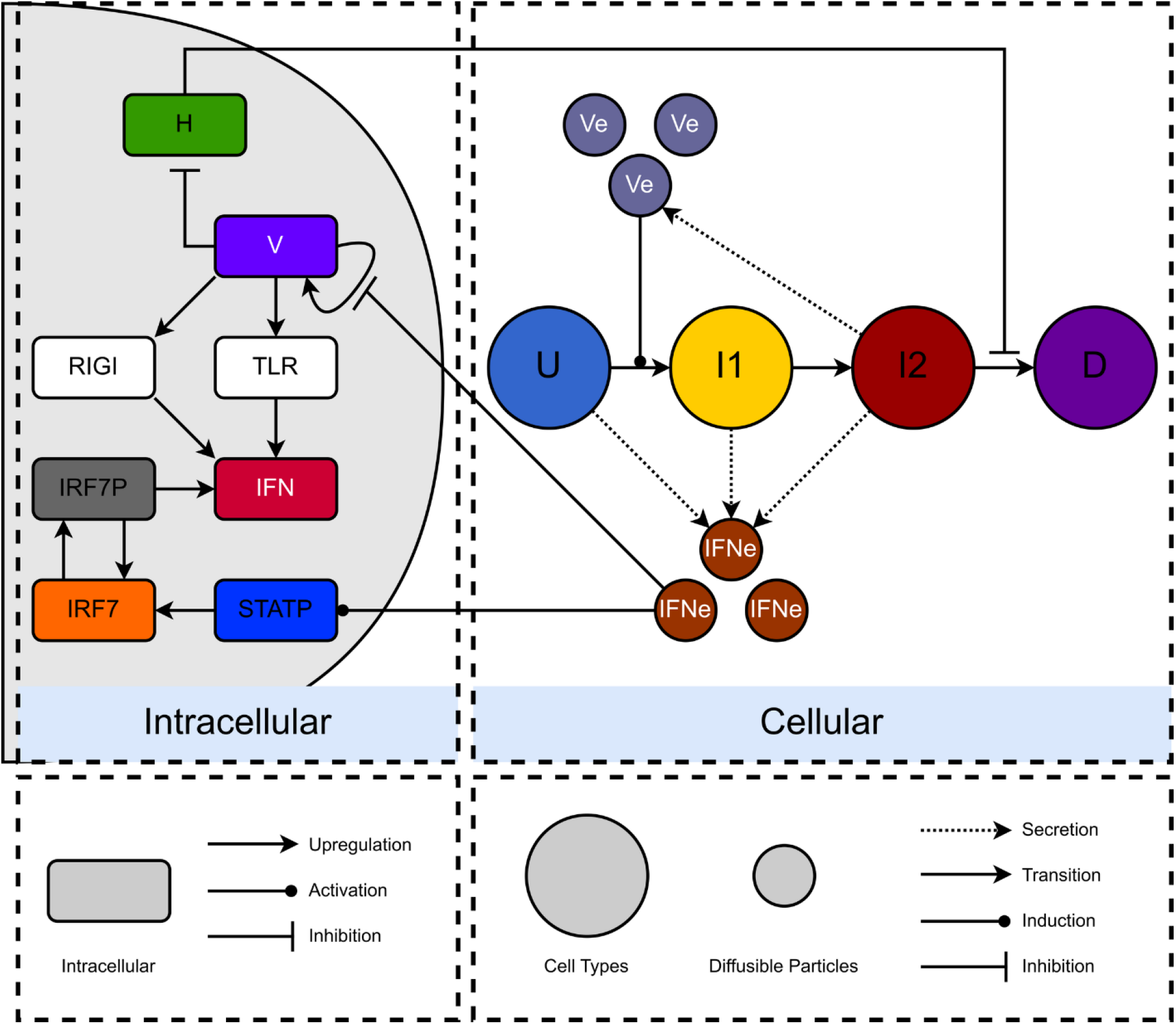
Conceptual Model Diagram. The model consists of an Intracellular sub model, which describes the innate immune response to infection, and the Cellular sub model, which defines changes in cell states and extracellular molecular diffusion. Uninfected cells (U, blue) can produce IFN via paracrine signaling. Eclipse phase cells (I1, yellow) produce IFN due to infection and paracrine signaling, and allow for virus replication but do not release virus. Virus releasing cells (I2, red) can produce IFN and release virus while dead (D, purple) cells do not interact with their surroundings. Cell state colors are conserved throughout this work. Each cell contains an instance of the ODE system, representing innate immune responses (RIG-I, TLR, IFN, IRF7, IRF7P, and STATP), viral infection, replication, and export (V), and cell viability (H). Type-I interferons (IFN_e_) and viral particles (V_e_) diffuse in the extracellular environment.

Cell State Transitions

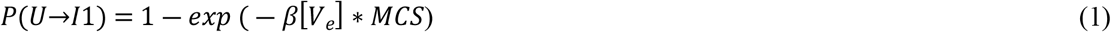

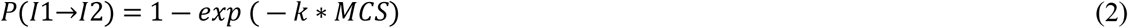

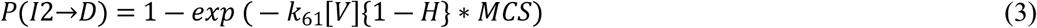

### Intracellular Model Equations and Rationale

Each cell within the simulation contains a set of differential equations to define intracellular virus replication, IFN response, and cell health. The IFN response is initiated when a cell becomes infected (as described by Eq. 1) and enters the eclipse phase, I1, where virus replication occurs but the export of new virions has not commenced (36). The intracellular virus’ exposed RNA is capable of inducing an IFN response during this phase via cytosolic retinoic acid inducible gene I (RIG-I) and cell surface toll-like receptor (TLR) family (37–40) sensor protein (Eq. 4, *k*_*11*_-*k*_*13*_) detection, which leads to the production of interferon α and β. Interferons are critical in activating the innate and adaptive immune responses including processes that suppress local virus replication. Here, interferon α and β are combined into a single representative molecule labeled interferon (IFN). Influenza’s nonstructural protein 1 (NS1) has strong antagonistic effects on multiple stages of the innate immune response (41,42), and near total antagonism of RIG-I. SARS-CoV’s NSp1 has been shown to exhibit similar RIG-I antagonism (43). Research shows an 87% conservation of NSp1’s genome between SARS-CoV and SARS-CoV-2 (13) and RIG-I antagonism in SARS-CoV-2. Thus, RIG-I’s activity was assumed zero for the baseline model.

Intracellular IFN (Eq. 4) is excreted to the external environment (Eq. 10), becoming environmental interferon, IFN_e_. This leads to autocrine and paracrine induction of more IFN and the activation of ISGs (44,45). The level of induction of ISGs is not modeled, but the effect of ISGs on inhibiting virus replication (46) is described by Eq. 9. IFN_e_ leads to the phosphorylation (activation) of STAT (indicated as STATP in Fig 1) through the JAK/STAT pathway, which in turn phosphorylates IRF7 to IRF7P and subsequently increases IFN production.

As the cell transitions from phase I1 to I2, viral export begins. The intracellular virus (V) is exported from the cell, becoming environmental virus, V_e_. The cell’s viral load impact on the cell’s health (H) is modeled with Eq. 8. As health nears zero, the likelihood of cell death increases dramatically (Eq. 3). When a cell is infected but resistant to infection, the virus may fail to sufficiently impact cell health and trigger cell death (Eqs. 8, 3). In this case, the cell will remain in the I2 state but will not produce additional virions.

Intracellular ODEs

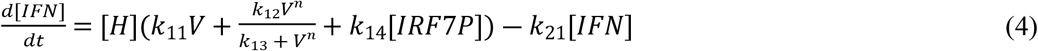

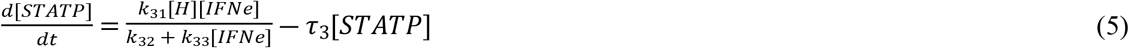

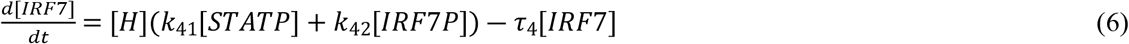

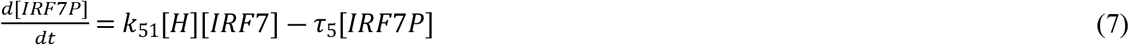

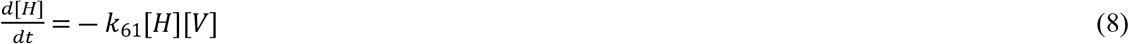

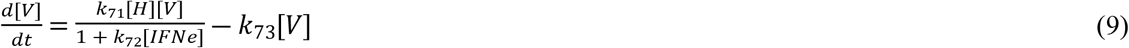

### Spatial Considerations: Diffusion of Interferon and Virus

The final component of the model is the diffusion of environmental interferon (IFN_e_) and virus (V_e_) across the simulated cell culture. Environmental interferon is excreted during the innate immune response by U, I1, and I2 cells, as described by Eq. 10. Environmental virus is exported by I2 cells, represented by Eq. 11. Both species diffuse across the cellular lattice and have first order decay rates. σ(*x,y,t*)is a spatial operator, indicating that the given species varies by cell (σ) located at lattice index *x,y* at time *t*. Each diffusive species is ready by the cell at lattice index *x,y* before undergoing diffusion at each timestep.

Extracellular Species

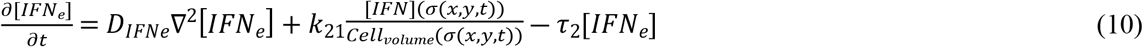

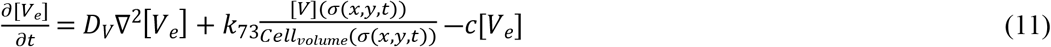

### Parameter Determination

Most modeled parameters were obtained from the lowest sum-of-squares error resulting from a previous parallel tempering Markov chain Monte Carlo parameterization of an ODE only model (33) to an infection of HBECs with wild-type A/Puerto Rico/8/1934 Influenza A (47). Additional parameters were estimated from results reported in literature (27,48–50). Table 1 gives a comprehensive list of model parameters and their origin. Virus diffusion values vary by 7 orders of magnitude over different media types (22), but the final value was chosen along with an IFN diffusion slightly lower (∼0.5 experimental value, 6 times simulated cell diameter) than experimental measurements (48,51) to securely place the baseline parameter set within in a regime of uncontained plaque growth. The cell state transition parameter β (34) was converted from TCID_50_ day^-1^ to PFU/mL day^-1^ for consistency with the IFN model’s prediction of viral load.

**Table 1.**
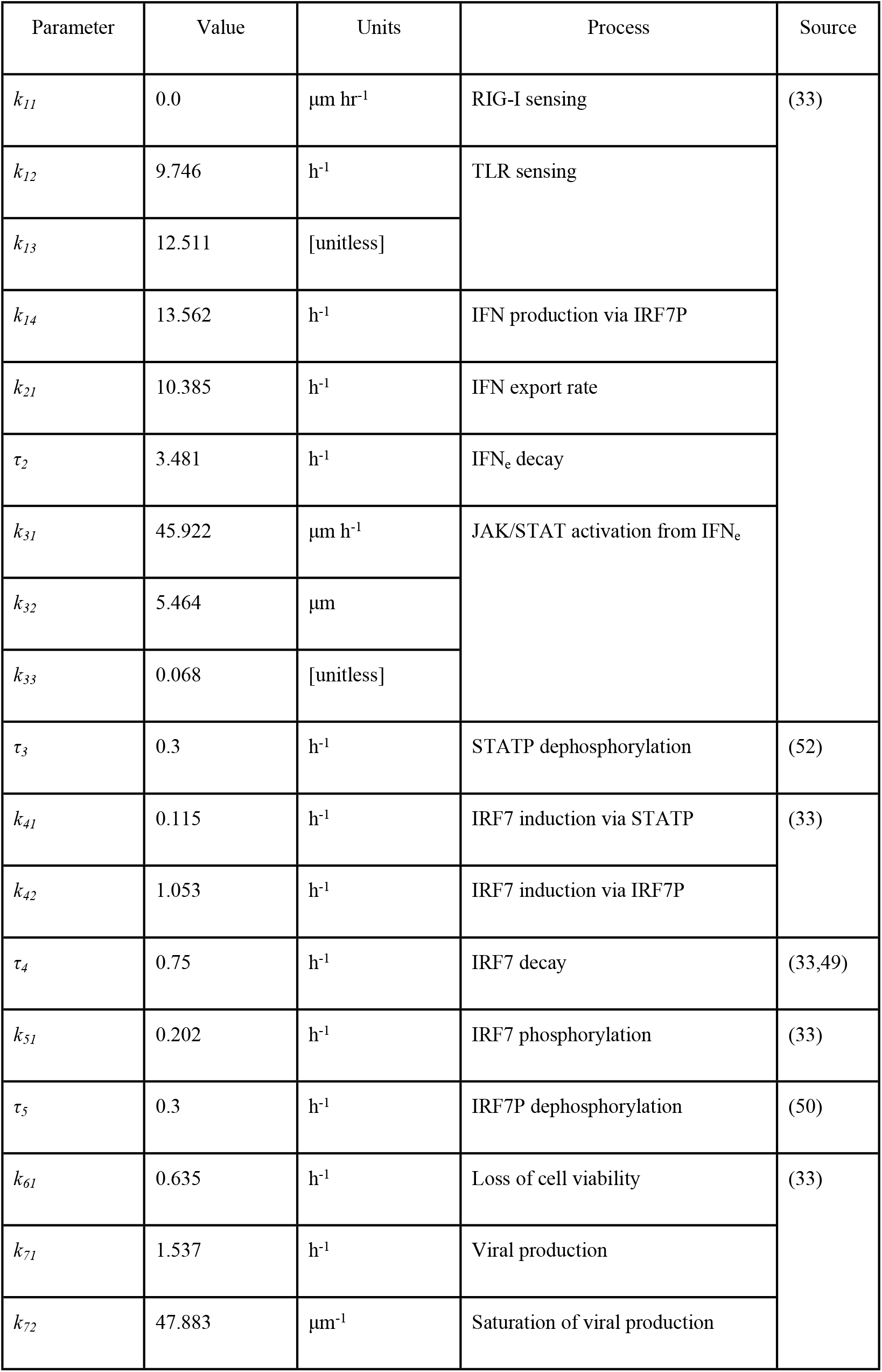
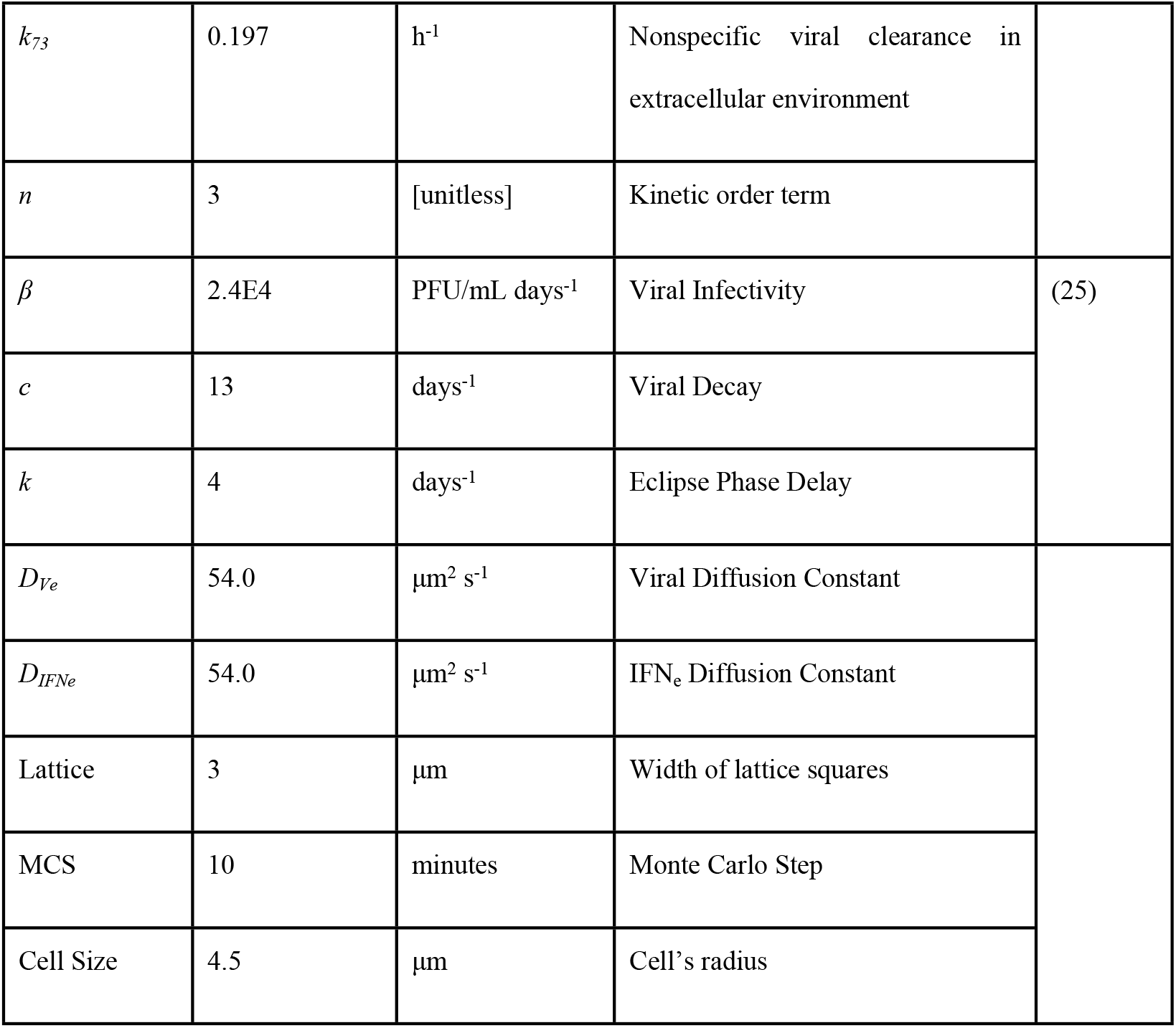
Baseline parameter values and sources.

### Plaque Growth Metrics

Viral plaques are visible areas of cell destruction which occur where a virus has spread across a cell culture. The plaque growth speeds of the eclipse phase (I1), virus releasing (I2), and dead (D) phases are used in several analyses. In the simulations, we determine these speeds by seeding a single infected cell in the center of a grid of cells and measuring the total area occupied by each cell state over time. We assume the plaque is circular, with the radius calculated to give plaque growth velocities. Simulations involve probabilistic infection events and stochastic cell transitions, however, the ultimate cell culture fates (*i*.*e*. plaque arrest versus uncontained growth) are deterministic. We averaged plaque growth metrics for sensitivity analyses over 20 simulations (S1 Text Fig 6). In experiments, plaque-plaque interference occurs when two or more plaques grow into the same spatial region, slowing the radial growth of all plaques involved. This paper simulates only growth of isolated plaques.

Plaque arrest occurs *in vitro* under specific conditions. Unlike models in which cells enter a virally resistant state at a fixed threshold of interferon, our model does not include an explicit resistant or immune state for the epithelial cells. Instead, the phenomenon of an antiviral state and plaque arrest arises naturally from the inclusion of ISG antagonism of viral replication in the ODE model (Eq. 6, *k*_*72*_). In our model, sufficiently high viral loads can overwhelm the cells’ antiviral state, and the resistant state can be lost over time as the cytokine signal decays. Virus releasing (I2) cells die at a rate depending on the cell’s internal viral load. Because the activated JAK/STAT pathway and ISGs inhibit virus replication, the viral load within individual simulated cells may not reach a level sufficient to trigger cell death.

## Results

### Multicellular Spatial Model of RNA Virus Infection and IFN Response Reproduces ODE Model Dynamics

As the spatial multiscale model presented here is predicated on an earlier ODE model, the multiscale model was first tested to see if it could reproduce similar dynamic responses to those obtained from the ODE system. Detailed in the methods, the ODE model was trained to data from HBECs (53) that were uniformly infected with an influenza virus at MOI = 5. The multiscale model was initialized under identical conditions to demonstrate that introduction of a stochastic cell state transition from virus releasing (I2) to dead (D) does not introduce differences between ODE and multiscale representations of the system. Despite homogenous starting conditions, a non-uniform cellular state field (Fig 2A,B) emerges due to the stochastic nature of cell transitions and the spatial patterns produced by IFN_e_ and extracellular virus. The average of each internal species over all cells is compared against the ODEs alone in Fig 2C.

**Fig 2.**
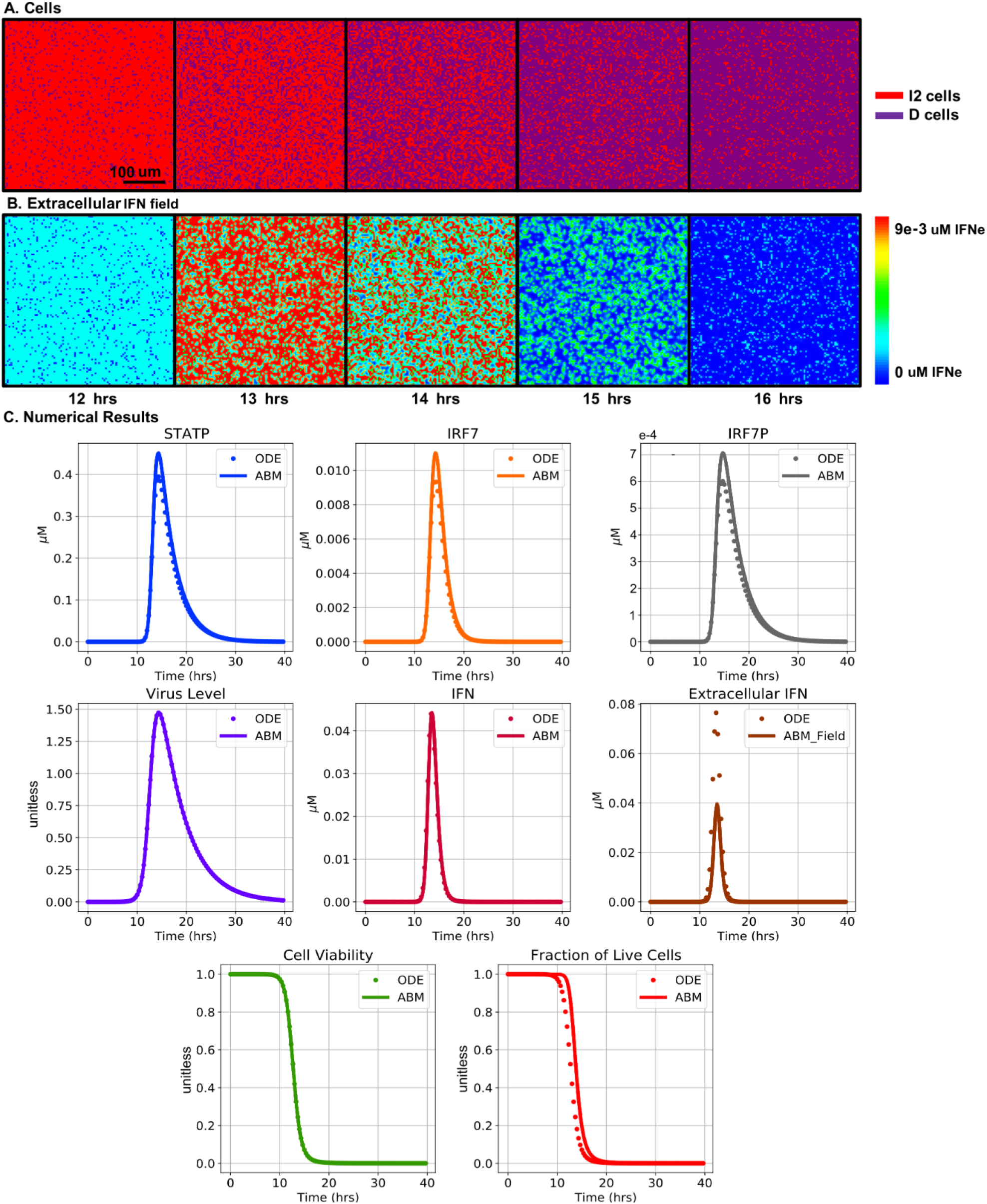
High MOI simulation and ODE comparison. All cells are initially infected with one viral particle, representing a multiplicity of infection of 5, as in the original data which the ODE system was trained to(53). The grid consists of 300×300 cells, representing approximately 2.7 x 2.7 mm of tissue. (A) Cells’ state (virus releasing (I2), red, and dead (D), purple). (B) Extracellular type-I interferon (high concentrations in red, low concentrations in blue). (C) Comparison between ODEs and spatial model. IFN_e_ per cell is the local concentration field which cells are exposed to. Note that differences arise due to a discrete number of cells rather than a normalized population value, from the stochasticity associated with cell transitions (Eq. 1-3), and diffusion of IFN and virus in the extracellular environment. Differences are minor due to the homogenous starting conditions and yield the same overall results as an ODE simulation. Simulations are averaged over 20 replicas (S1 Text Fig 6).

The average concentrations of intracellular species and viral titer are like those of the ODE only model. Living cell fraction dynamics are delayed in the multicellular spatial model, as the transition from I2 to D is now discrete (whole cells) and stochastic (probability based transitions), rather than continuous (normalized population) and deterministic (ODE rate of death). Under these homogeneous, high MOI starting conditions, the multicellular spatial model, under both a diffusive regime and a perfectly mixed system (data not shown), tracks the ODE model very well. The system is less sensitive to spatial differences under these starting conditions, since all cells are initially infected, fewer stochastic cell state transitions are possible, and complete cell death occurs much earlier than in low MOI scenarios. Since the multiscale model can reproduce ODE system dynamics under well mixed experimental conditions despite the differences in how the models function, the multiscale model serves as a foundation for exploration of the system under novel conditions: an exploration of low MOI conditions and spatially heterogeneous starting conditions which are highly relevant for *in vivo* organism infections and *in vitro* plaque growth assays.

### Multicellular Spatial Model Recapitulates Plaque Formation and Growth Dynamics

High MOI, or single cycle experiments, are useful for maximal viral titer experiments and determining cell survival time, but they do not provide information about viral spread and spatial cytokine response patterns, do not allow for studying viral growth arrest, and are not representative of the conditions of a normal infection. We wished to explore low MOI, or multicycle growth experiments, *in silico*. This section serves as an exploration of plaque growth assays, a low MOI experimental technique that was recreated in the multiscale model. A plaque is a visible area of dead or injured cells that form in cell culture and in the lung of infected animal models as the virus spreads from a point of infection. Fig 3 (left) shows multiple plaques in a culture of cells infected with A/Miss/2001 H1N1 influenza virus. To replicate plaque formation *in silico*, we simulated an experiment starting with a single A/Miss/2001 H1N1 infected cell seeded at the center of a cell culture. Fig 3 demonstrates that the multicellular spatial model can capture the formation of a viral plaque observed *in vitro*.

**Fig 3.**
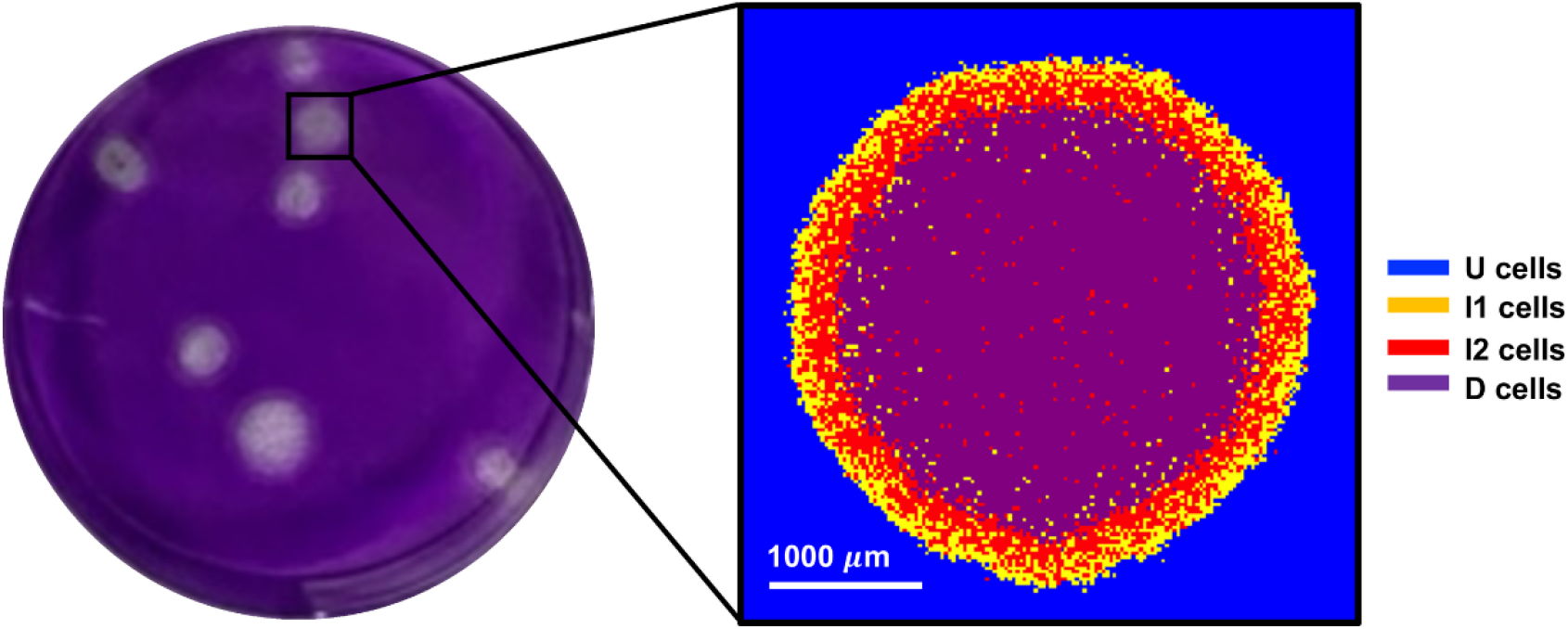
A comparison of an experimentally observed plaque resulting from cells infected with influenza A/Miss/2001 (H1N1) and multicellular spatial plaque simulation at 48 hours. Both plaque and simulation seeded at a 2E-5 MOI (21). Next, we explored the model’s plaque growth dynamics. Fig 4A, B show experimentally observed plaque growth (data reproduced from (22)). While the increase in the number of viral particles during infection is typically exponential, plaque growth, as measured by the change in the diameter of the plaque over time, is linear. There are two experimentally defined boundaries of the plaque, namely, the outer edge of the dead cells (equivalent to D in the model) and the outer edge of the infected cells (equivalent to I1 in the model). The model distinguishes eclipse phase (I1) and virus releasing (I2) cells, which is not normally seen in experimental plaque growth assays. Seeding the simulation with a single initially infected cell, Fig 4C shows that the multicellular spatial model replicates these experimental observations. A lag phase during which the initially transfected virus is establishing its replication and export is seen both experimentally (Fig 4A,B) and in the simulation (Fig 4C). Fig 4D-F shows virus spread, plaque formation and IFN diffusion over time. Fig 4G provides cell state trajectories. There is a lag time of 20 hours until dead cells are present, after which the growth in the radius of the dead cells increases linearly. The growth rate of the plaque remains constant until host cells are depleted. A temporary decrease in environmental virions (Fig 4H) and total IFN concentration in the culture (Fig 4I) is exhibited at 18 hours, during the transition from initially infected cells (I1) to subsequently infected (I2) cells. Neighboring cells are infected starting at 8 hours, and transition to virus releasing cells starting at 10 hours. During this time, the initially infected cell is dead and no longer exporting viral particles, while extracellular virus is decaying.

In summary, our multicellular spatial model of epithelial infection and IFN response recapitulates plaque growth dynamics seen *in vitro* using the parameters from the source ODE model. The model’s ability to simulate high and low MOI experiments and reproduce phenomena seen experimentally, without any training to these conditions, gives confidence in its predictive capabilities in novel circumstances. The next five sections of Results are based on simulations of plaque growth assays to give spatialized insights into the mechanisms regulating plaque growth, an element missing from ODE and high MOI research.

### Increased STAT Activity Leads to Arrested Plaque Growth and Reduces Final Plaque Diameter

The JAK/STAT pathway triggers an immune reaction via paracrine signaling facilitated by cytokines. This mechanism has been implicated in improved H1N1 influenza survival in a murine model (54), and more broadly in host response to SSRNA viruses such as Japanese Encephalitis Virus (55). To assess the impact of STATP activity on plaque growth dynamics, we simulated plaque growth values of *k*_*31*_ (Eq. 2, Table 1), which represents the ability of extracellular interferons to activate the JAK/STAT pathway, from its baseline value (45.9 μm hr^-1^, Table 1) by 10 and 100 fold were carried out. For each value of *k*_*31*,_ we show the plaque at 80 hours post infection (Fig 5A) and the cell state dynamics over time (Fig 5B). At baseline, unconstrained plaque growth is observed. Values of *k*_*31*_ *≥*459.22 μm hr^-1^ (10 times baseline) lead to arrest of plaque growth. Increasing *k*_*31*_ *≥* 4592.2 μm hr^-1^ (100 times its baseline value) reduces the time it takes to reach plaque growth arrest, resulting in smaller plaques. In all, these results show that modulating STATP activity within the multiscale model can arrest plaque growth and smaller final plaque size.

The model asserts that a change in *k*_*31*_ impacts plaque growth rate, as quantified by the rate of change of the plaque at the end of the simulation (Fig 5C). The area under the curve (AUC) of extracellular virus (Fig 5D) and environmental IFN produced (Fig 5E) show reduced infection success and a stronger immune response for increasing *k*_*31*_. Values of *k*_*31*_ *≥* 4592.2 μm hr^-1^ lead to non-physiological unbounded production of IFN, due to the lack of an IFN-mediated cell death mechanism in the model. This limits predictions to qualitative observations, rather than quantitative predictions of viral titer or IFN_e_ concentration. Plaque metrics (Fig 5C-E) demonstrate the magnitude of the immune response and quantify the decrease in overall viral production.

### Elevated RIG-I Activity Delays Cell Death and Increases IFN

Greater viral inhibition of RIG-I signaling via nonstructural protein production often increases viral infection severity (41–43). We investigated the effects of varying the antagonistic strength on plaque growth dynamics *in silico*. In all simulations thus far, RIG-I is assumed to be fully antagonized by the invading virus (*k*_*11*_ = 0). We ran single plaque simulations with 0%, 25%, 50%, 75%, and 100% RIG-I activity relative to the value in an NS1 knockout training simulation using A/Puerto Rico/8/1934 (dNS1PR8). At 80 hours post infection (Fig 6A) plaque radius is nearly unchanged in all cases. However, the cell type composition (Fig 6B) changes significantly. Higher levels of RIG-I signaling (larger values of *k*_11_) lead to less cell death (Fig 6C), lower viral titer (Fig 6D), and much greater interferon production (Fig 6E).

Final plaque radii remain almost constant as RIG-I activity is varied from 0 to 1. Increasing RIG-I activity over baseline levels delays cell death. This effect originates from greater interferon responses, and in turn, lower viral titers and an increasing occurrence of failed infections which do not lead to significant viral replication or export. We predict that enhancing RIG-I activity and/or reducing nonstructural protein production in an experiment should reduce viral growth by allowing greater cell survival time in which the innate immune response can further propagate.

### Interferon Prestimulation Arrests Plaque Growth

In experiments, prestimulation of cell cultures with type-I interferons, protects against SARS-CoV (13), SARS-CoV-2 (14), and influenza (15). We simulated an infection with such prestimulation by exposing uninfected cells to IFN_e_, after which a single cell was infected to initiate the plaque. We found that prestimulation under these conditions entirely arrests plaque growth; while the same initial infection in a field of naive cells under identical simulation conditions would result in the infection and eventual death of all simulated cells (Fig 4 C,G). Fig 7A shows that IFN-prestimulation slows or arrests plaque growth. Only the initially infected cell dies, while the proportion of eclipse phase (I1) cells steadily decreases after 20 hours, indicating a cessation of new infections (Fig 7B). The extracellular viral load (Fig 7C) also decreases after 20 hours. Extracellular IFN (Fig 7D) indicates an elevated cytokine profile from the prestimulated cells.

**Fig 4.**
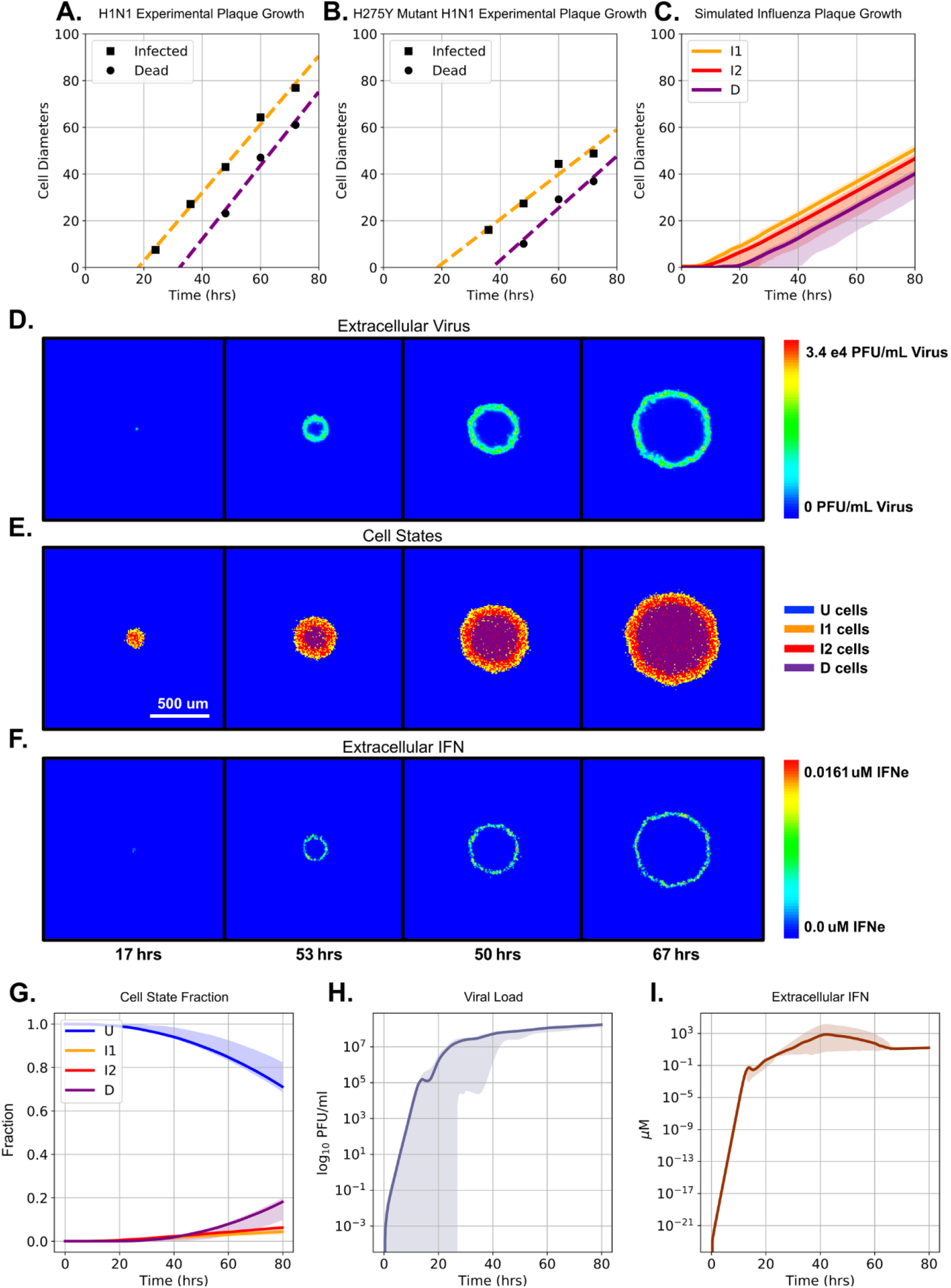
Single Plaque Growth Simulations Recapitulate Experimental Dynamics. (A, B) Radial growth of wild-type and H275Y mutant A/Miss/3/2001 (HIN1) plaques, respectively. Data reconstructed from (22). Squares indicate outer edge (infectious cells) and circles indicate inner edge (dead cells) of the plaques. Dotted lines show a linear regression for visualization of plaque growth rate, not a model fit. (C) Simulated plaque growth without any training recapitulates the lag phase and linear growth of the experimental plaques. Radial growth is determined as described in the Methods. The radius is averaged over 20 simulations (S1 Text Fig 6) and the shaded areas indicate the maximum and minimum observed values. Panels D, E and F show images of simulated plaque growth, the cell states and the extracellular IFN, respectively, over time, for one instance of seeding a single infected cell in the center of the cell lattice. Time progresses from left to right. Panels G, H and I show the mean (solid line) and max/minimum (shaded areas) of the simulated cell states, viral load and extracellular IFN titer, respectively, observed over 20 simulations (S1 Text Fig 6).

**Fig 5.**
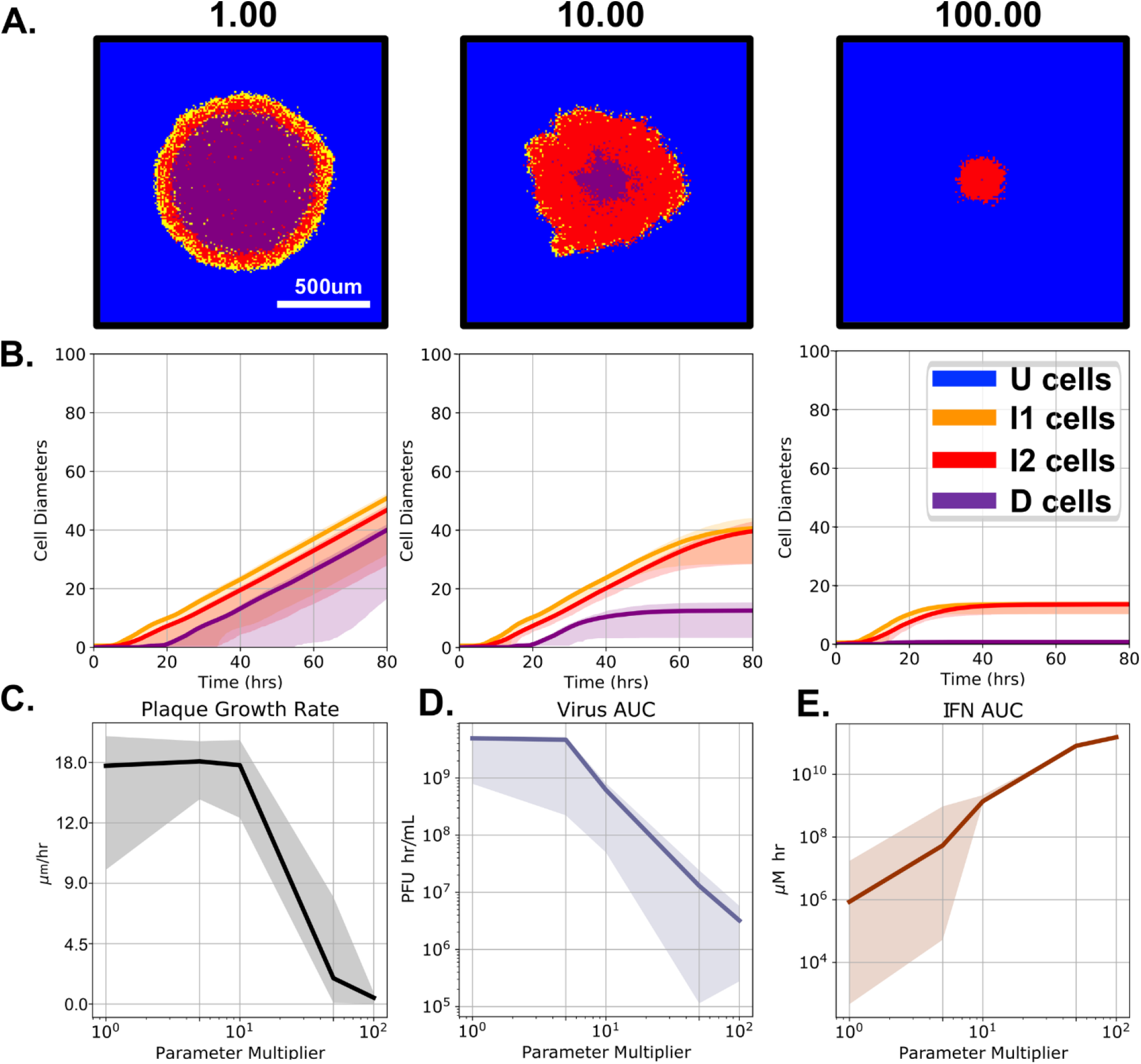
Elevated STATP activity (*k*_*31*_) leads to arrested plaque growth. (A) Images of the plaque at 80 hours post infection for a single simulation and (B) the mean (solid line) and max/minimum (shaded regions) over 20 simulations of the plaque radial growth over time for when *k*_*31*_ is 1x, 10x, or 100x larger than its nominal value. We also show (C) plaque growth rate at 80 hours, (D) area under the curve (AUC) of the virus concentration, and (E) AUC of the IFN concentration for different values of *k*_*31*_. Shaded areas represent the minimum and maximum values over 20 replicates.

**Fig 6.**
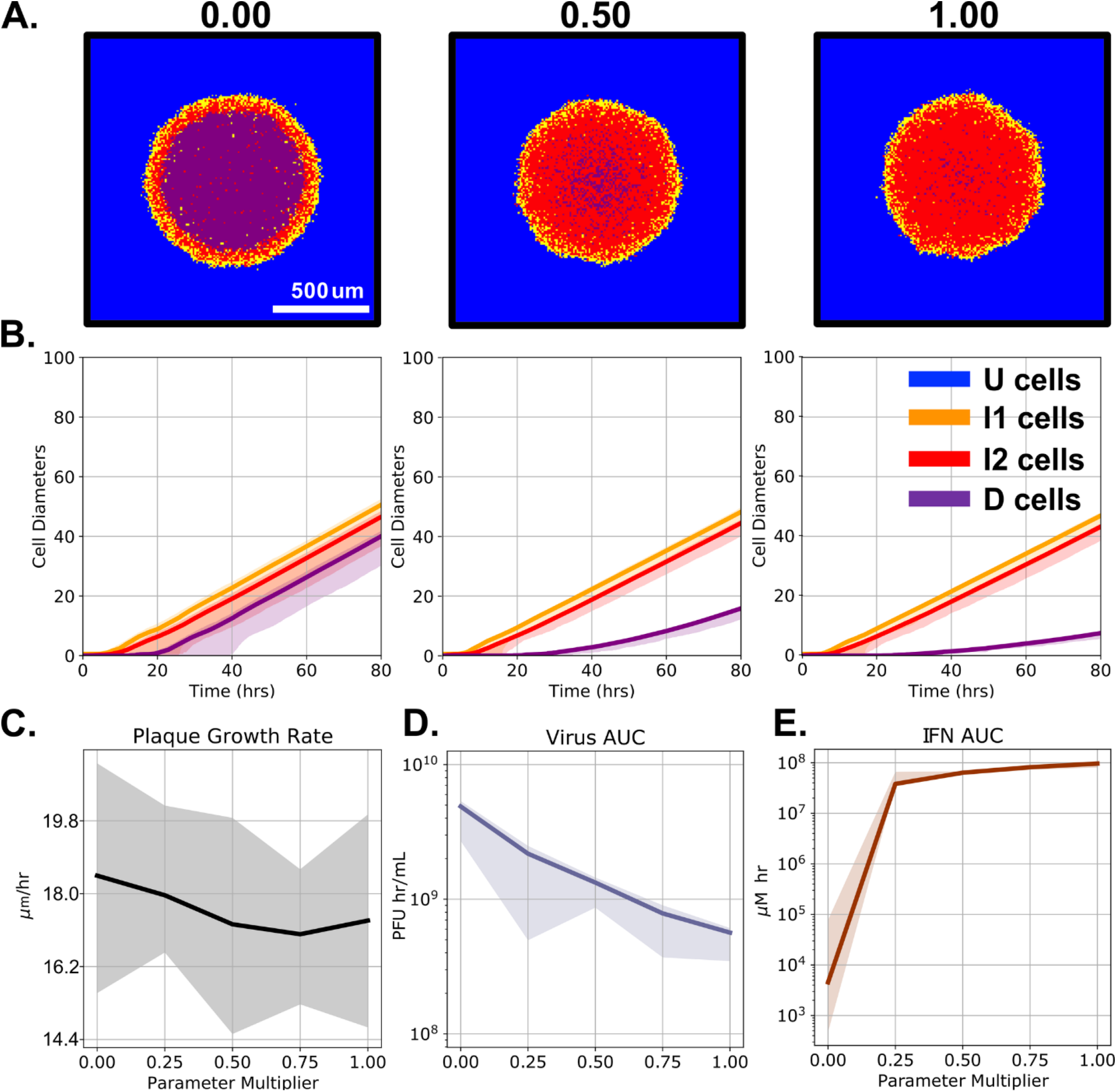
Reduced viral RIG-I antagonism lowers plaque growth rates and viral titers, slows cell death, and increases interferon production. (A) Images of plaques at 80 hours post infection for a single simulation and (B) the mean (solid line) and max/minimum (shaded regions) over 20 simulations of the plaque radial growth over time for when *k*_*11*_ is 0x (0, Table 1), 0.5x (5e5), or 1x (10e5) its nominal value in an NS1 knockout strain. We also show (C) the plaque growth rate at 80 hours, (D) area under the curve (AUC) of viral load, and (E) AUC of IFN concentration for different values of *k*_*11*_. Shaded areas represent the minimum and maximum values over 20 replicates.

**Fig 7.**
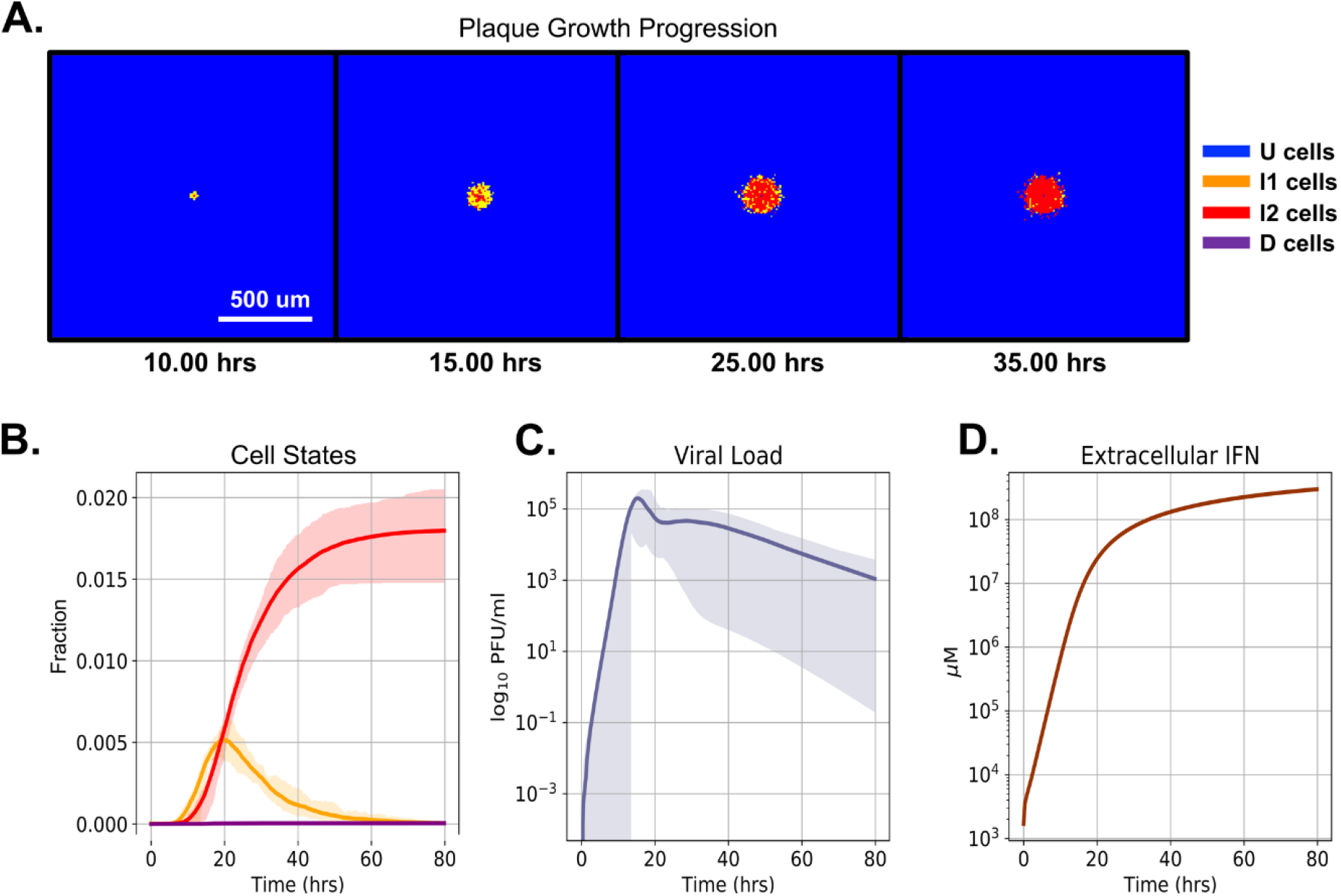
Prestimulating cells with type-I interferon leads to plaque growth arrest in simulations. The cells are simulated as washed with extracellular IFN, which is removed immediately before infection. (A) Time series of plaque growth for a representative simulation. The cell states (B), extracellular viral load (C) and concentration of extracellular IFN (D) correlate with plaque progression and severity. The solid line indicates the mean and shaded areas represent the minimum and maximum over 20 replicates.

After prestimulation with IFN, infected cells survive because the viral load remains low, minimizing damage to the infected cell’s health, which remains above the threshold to induce cell death. The persistence of virus releasing cells does not affect whether the plaques arrest or continue to grow, as these cells stop producing virions. This condition is akin to a failed infection event, wherein the ISG effects would prevent the viral replication from occurring.

### Relative Diffusion Constants of Interferon and Virus Determine Plaque Growth Dynamics

We explored increased interferon diffusion constants, which emulate different media types *in vitro* (22) or dendritic cell signal amplification *in vivo* (56). We performed a two dimensional parameter variation of virus and interferon diffusion coefficients. The interferon diffusion coefficient was varied from 54 um^2^ s^-1^ to 2160 um^2^s^-1^ (1.0 to 40.0 times the baseline diffusion constant) and the virus diffusion constant from 54 um^2^s^-1^ to 216 um^2^s^-1^ (1.0 to 4.0 times the baseline diffusion constant). We evaluated the outcome of the simulation by averaging the growth rate of the plaque at the end of the simulation over 20 simulation replicates. If the average growth rate was 0 we classified the parameters as leading to plaque arrest, otherwise we classified the parameters as leading to continued growth (blue). In our simulations an interferon diffusion constant of about 8- to 10-fold times the viral diffusion constant led to plaque growth arrest (Fig 8). The nonlinear boundary between the domains suggests that viral diffusion eventually ceases to be the rate limiting factor in plaque growth and that higher interferon diffusion constants do not saturate the cells’ ability to respond to the infection within the scales investigated. Larger interferon diffusion constants lead to a broader area of protection around an IFN signaling cell, but this protection can be overcome by a greater viral particle diffusion constant.

**Fig 8.**
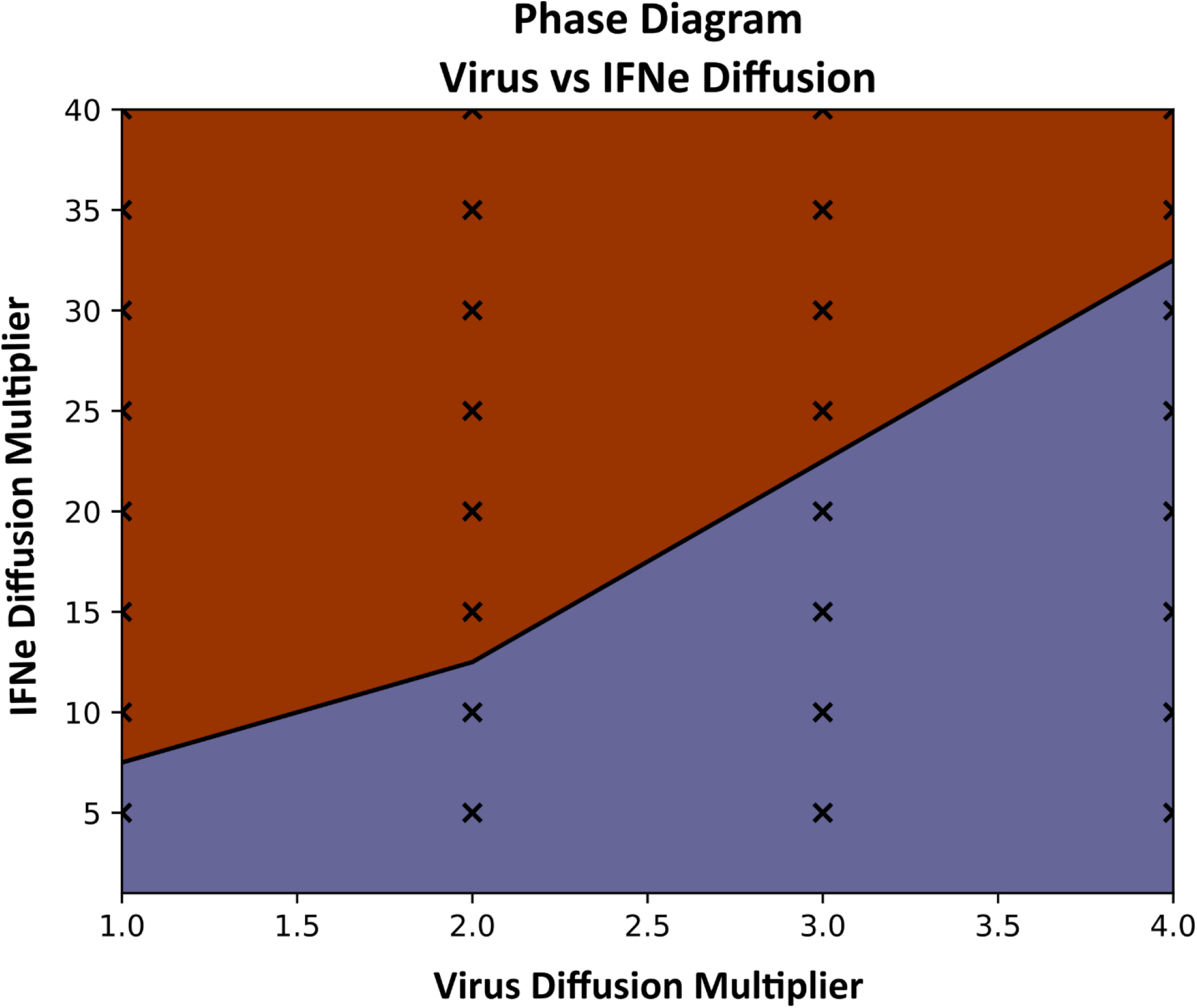
Dependence of plaque arrest on viral and INF diffusion constants. In the orange (top) region plaques arrest by 80 hours. In the purple (bottom) region, plaques continue to grow. Crosses show simulated parameter combinations. S1 Text Fig 2 shows individual simulations’ cell state progression.

### Sensitivity Analysis Reveals Changing Controlling Parameters Between Regimes

To determine how individual parameters affect the growth of plaques we performed local sensitivity analyses for three regimes in parameter space; the baseline parameter set (Table 1), an elevated JAK/STAT signaling regime (*k*_*31*_ = 688.5 μm hr^-1^, 15 times baseline), and an elevated interferon diffusion regime (D_IFNe_ = 540.0 μm^2^ s^-1^, 10 times baseline). For each regime, we ran 20 simulations using the regime’s nominal parameter values. Then, we perturbed each parameter individually ±25%, computed 20 simulations for each new parameter set, and performed statistical analysis on several sensitivity metrics derived from the simulated trajectories. The sensitivity metrics include the percent change from the average of the nominal simulation of the plaque growth rate, the extracellular virus and IFN maxima and AUC of the extracellular virus and IFN. Statistical significance of the change in each metric from their nominal values was determined using a Student’s t-test. Values are reported in the Supporting Information (S1 Text Figs. 7-9). Increasing and decreasing perturbations to the parameter values primarily led to directionally consistent changes in the sensitivity metrics (*e*.*g*. if the metric value increased when the parameter was changed by +25%, then the metric value also decreased when the parameter value was changed by -25%). The sensitivity metrics in Fig 9 average the absolute values of the metric for increased and decreased parameters. The top row of Fig 9 shows the cell state progression of a plaque growth assay under each regime.

**Fig. 9.**
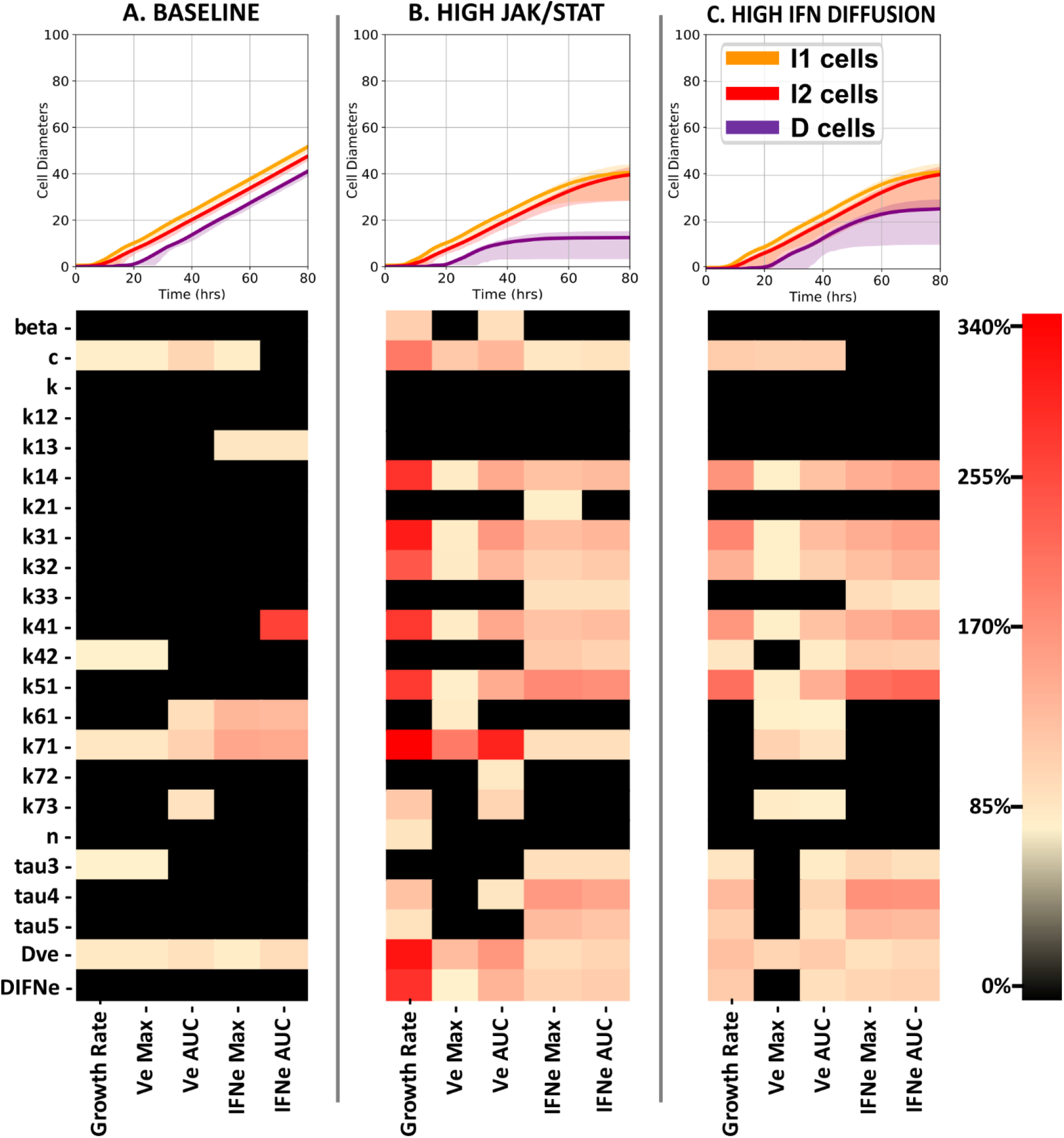
Local sensitivity analysis under three simulation regimes. A. “Baseline” corresponds to the baseline parameters in Table 1. B. “High JAK/STAT” corresponds to a 15-fold increase in the phosphorylation rate of STATP via the JAK/STAT pathway (parameter *k*_*31*_), with all other parameters as in Table 1. C. “High IFN Diffusion” corresponds to a 10-fold increase in the diffusion constant of IFN_e_, with all other parameters as in Table 1. Sensitivity analyses varied each parameter one at a time ± 25% around its baseline value and quantifying the average plaque growth rate at the end of the simulation, the maximum extracellular virus (V_e_) and interferon (IFN_e_) levels, and the area under the curve (AUC) for both V_e_ and IFN_e_.

Previous sections demonstrated that variation in multiple parameters could lead to either continuous or arrested plaque growth. The baseline parameter set leads to continuous growth of the plaque. In this regime, STATP’s dephosphorylation (*τ*_*3*_) and induction of IRF7 (*k*_*41*_, *k*_*42*_) and the virus’ rate of production (*k*_*71*_), diffusion (*D*_*Ve*_), and nonspecific environmental clearance (*c*) have large effects on outcomes of the simulation. For example, increasing the viral replication rate and virus diffusion constant lead to faster plaque growth, whereas increasing the virus decay rate in the extracellular environment slows plaque growth. The High JAK/STAT and High IFN Diffusion regimes have arrested plaque growth. In these regimes, the many parameters associated with the activation of paracrine signaling have substantial effects, with higher sensitivity in the High JAK/STAT regime than in the High IFN Diffusion regime. Diffusion constants further from the boundary between the plaque growth and arrested growth regimes, have greater sensitivity to virus associated parameters and weaker sensitivity to IFN associated parameters.

## Discussion

We have explored the spatiotemporal behavior of a multicellular spatial model of the cellular innate immune response to viral infection. To do so, we developed a general approach to adapting an existing ODE model of spatially homogenized dynamics to create a calibrated spatiotemporal model, which has the potential to enable the addition of spatial investigations to many existing ODE frameworks in systems biology.

One of the striking takeaways is the bifurcation between plaques growing without limit and plaque arrest. This insight into the bifurcation between uncontained plaque growth and plaque arrest would be impossible without a spatial aspect present in the model and can be readily understood as a metric of disease progression and severity. We demonstrated plaque arrest under elevated interferon diffusion (Fig 8), elevated paracrine signaling (Fig 5), and interferon prestimulation (Fig 7).

This model demonstrates the dramatic dependence of cell culture fate on interferon and virus diffusion, shown in Fig 8. Increased interferon diffusion results in a limited viral plaque by initiating IFN production and ISG effects in a larger local neighborhood of cells. The resulting antiviral state is an emergent property of the interactions between multiple biological scales: spatial propagation of infection and cytokine signal in the extracellular environment and intracellular transduction of IFN signaling, rather than the traditional transition of cells to a distinct virally resistant state. Our model consistently predicts a low viral titer in the dead cell region of plaques, shown in Fig 4D. This could be explained as an evolutionary strategic adaptation; infectious particles in this region do not contribute to the replication of the virus. Instead, viral particles with high enough diffusion only to reach new hosts, both at the tissue level and in the epidemiological sense, maximize the reproductive success of the virus given limited host resources for replication. In response, the hosts’ production of signaling cytokines (Fig 4F) are concentrated primarily near the site of infected cells which are actively producing virus, attracting immune cells to the wave front of infection via chemotaxis rather than allocating resources to the cellular debris in the center of the plaque. *In vivo*, virus and IFN don’t diffuse passively, due to cilia action, periciliary fluid, and mucosal layer driven advection. We will explore these active transport mechanisms in future work.

Elevating STATP activity, as affected by *k*_*31*_ and explored in Fig 5, can lead to non-physiologically high IFN production. This case can lead to cells remaining in the I2 state without producing additional virions. Biologically, these cells may be reinfected or undergo cytokine-induced programmed cell death, but these outcomes were not within the scope of the model presented here. However, which of the two outcomes is reached by the simulation, *i*.*e*. arrest or continued growth of plaques, is unaffected and qualitative observations can still be made regardless of the unbounded IFN production. In future work, we will consider mechanisms to cut off IFN production in a physiologically meaningful way and model immune cells and drug interactions.

Our IFN pretreatment results can explain early clinical results which show inhaled interferon β as a promising treatment for COVID-19 (57,58). A tenfold increase in interferon production over the baseline parameters of the model is within biological possibility. Our model is based on data from HBECs, which have a limited capacity for cytokine signaling comparable to *in vivo* production from infected epithelial tissue. The recruitment of dendritic cells and their resulting signal amplification (59) *in vivo* in the local area of infection could create regions of elevated IFN and cells in an antiviral state induced by ISGs, and the clearance of infected cells and free viral particles by other immune cell classes would limit viral spread compared to a culture of purely HBECs.

The multicellular model presented here serves as a jumping off point for integration with immune cell action and other aspects of the body’s response to infection, as the spatial cytokine information could be used for chemotaxis and adaptive immune cell responses. It is our hope to carry these out collaboratively, with the goal of understanding severe SARS-CoV-2 infections and accelerating treatment development by simulating interventional strategies. Targeted experiments can bolster model predictions, and in turn inform experiments in a positive feedback loop between clinical data, experiment, and computational work. Finally, the methodology of creating data calibrated multiscale models in CompuCell3D based on existing ODE or rule-based systems holds enormous potential for investigating the spatial aspect of biological systems.

## Acknowledgments

We would like to thank Dr. John Alcorn, Dr. Douglas S. Reed and Dr. Katherine J. O’Malley at the Department of Immunology and University of Pittsburgh Center for Vaccine Research for providing the plaque assay image used in Fig 3. We would also like to thank Emily E. Ackerman for her help in editing the manuscript. This research was supported in part by Lilly Endowment, Inc., through its support for the Indiana University Pervasive Technology Institute.

## Supporting Information Captions

**S1 Text. Supporting Information Figures.**

## References

1. WHO. Global Influenza Strategy 2019-2030.

2. THE GEOGRAPHY AND MORTALITY OF THE 1918 INFLUENZA PANDEMIC on JSTOR [Internet]. [cited 2020 Nov 30]. Available from: https://www.jstor.org/stable/44447656?seq=1#metadata_info_tab_contents

3. Weekly epidemiological update -16 February 2021 [Internet]. [cited 2021 Feb 22]. Available from: https://www.who.int/publications/m/item/weekly-epidemiological-update16-february-2021

4. Yang Y, Tang H. Aberrant coagulation causes a hyper-inflammatory response in severe influenza pneumonia. Cell Mol Immunol. 2016;13(4):432–42.

5. Amor S, Fernández Blanco L, Baker D. Innate immunity during SARS‐CoV‐2: evasion strategies and activation trigger hypoxia and vascular damage. Clin Exp Immunol [Internet]. 2020 Nov 12 [cited 2020 Dec 4];202(2):193–209. Available from: https://onlinelibrary.wiley.com/doi/10.1111/cei.13523

6. Shibabaw T, Molla MD, Teferi B, Ayelign B. Role of ifn and complements system: Innate immunity in sars-cov-2 [Internet]. Vol. 13, Journal of Inflammation Research. Dove Medical Press Ltd; 2020 [cited 2020 Dec 4]. p. 507–18. Available from: /pmc/articles/PMC7490109/?report=abstract

7. AbdelMassih AF, Ramzy D, Nathan L, Aziz S, Ashraf M, Youssef NH, et al. Possible molecular and paracrine involvement underlying the pathogenesis of COVID-19 cardiovascular complications. Cardiovasc Endocrinol Metab [Internet]. 2020 [cited 2020 May 11];Publish Ah:1–4. Available from: https://journals.lww.com/cardiovascularendocrinology/Fulltext/9000/Possible_molecular_and_paracrine_involvement.99948.aspxNS-

8. Baskin CR, Bielefeldt-Ohmann H, Tumpey TM, Sabourin PJ, Long JP, García-Sastre A, et al. Early and sustained innate immune response defines pathology and death in nonhuman primates infected by highly pathogenic influenza virus. Proc Natl Acad Sci U S A [Internet]. 2009 Mar 3 [cited 2020 Dec 4];106(9):3455–60. Available from: http://www.pnas.org/cgi/content/full/

9. Kobasa D, Jones SM, Shinya K, Kash JC, Copps J, Ebihara H, et al. Aberrant innate immune response in lethal infection of macaques with the 1918 influenza virus. Nature. 2007;445(7125):319–23.

10. Cilloniz C, Pantin-Jackwood MJ, Ni C, Goodman AG, Peng X, Proll SC, et al. Lethal Dissemination of H5N1 Influenza Virus Is Associated with Dysregulation of Inflammation and Lipoxin Signaling in a Mouse Model of Infection. J Virol [Internet]. 2010 Aug 1 [cited 2019 Sep 24];84(15):7613–24. Available from: http://www.ncbi.nlm.nih.gov/pubmed/20504916

11. Peiris JSM, Cheung CY, Leung CYH, Nicholls JM. Innate immune responses to influenza A H5N1: friend or foe? Trends Immunol [Internet]. 2009 Dec [cited 2019 Sep 23];30(12):574–84. Available from: http://www.ncbi.nlm.nih.gov/pubmed/19864182

12. Shinya K, Okamura T, Sueta S, Kasai N, Tanaka M, Ginting TE, et al. Toll-like receptor pre-stimulation protects mice against lethal infection with highly pathogenic influenza viruses. Virol J. 2011;8:97.

13. Lokugamage K, Hage A, de Vries M, Valero-Jimenez A, Schindewolf C, Dittmann M, et al. Type I interferon susceptibility distinguishes SARS-CoV-2 from SARS-CoV. bioRxiv Prepr Serv Biol [Internet]. 2020 [cited 2020 Aug 5]; Available from: /pmc/articles/PMC7239075/?report=abstract

14. Lokugamage KG, Hage A, Schindewolf C, Rajsbaum R, Menachery VD. SARS-CoV-2 is sensitive to type I interferon pretreatment. bioRxiv Prepr Serv Biol [Internet]. 2020 [cited 2020 Aug 4];179:104811. Available from: https://doi.org/10.1101/2020.03.07.982264

15. Qiao L, Phipps-Yonas H, Hartmann B, Moran TM, Sealfon SC, Hayot F. Immune response modeling of interferon beta-pretreated influenza virus-infected human dendritic cells. Biophys J [Internet]. 2010 Feb 17 [cited 2019 Sep 9];98(4):505–14. Available from: http://www.ncbi.nlm.nih.gov/pubmed/20159146

16. Zhou Q, Chen V, Shannon CP, Wei X-S, Xiang X, Wang X, et al. Interferon-α2b Treatment for COVID-19. Front Immunol [Internet]. 2020 May 15 [cited 2020 Sep 18];11:1061. Available from: https://www.frontiersin.org/article/10.3389/fimmu.2020.01061/full

17. Covid: Large trial of new treatment begins in UK -BBC News [Internet]. [cited 2021 Jan 15]. Available from: https://www.bbc.com/news/health-55639096

18. Smith AM. Validated models of immune response to virus infection. Vol. 12, Current Opinion in Systems Biology. Elsevier Ltd; 2018. p. 46–52.

19. Gregg RW, Sarkar SN, Shoemaker JE. Mathematical modeling of the cGAS pathway reveals robustness of DNA sensing to TREX1 feedback. J Theor Biol [Internet]. 2019 Feb 7 [cited 2019 Jun 28];462:148–57. Available from: https://www.sciencedirect.com/science/article/pii/S0022519318305484

20. Pawelek KA, Huynh GT, Quinlivan M, Cullinane A, Rong L, Perelson AS. Modeling Within-Host Dynamics of Influenza Virus Infection Including Immune Responses. Antia R, editor. PLoS Comput Biol [Internet]. 2012 Jun 28 [cited 2020 Dec 15];8(6):e1002588. Available from: https://dx.plos.org/10.1371/journal.pcbi.1002588

21. Bocharov GA, Romanyukha AA. Mathematical Model of Antiviral Immune Response III. Influenza A Virus Infection. J Theor Biol [Internet]. 1994 Apr [cited 2019 May 30];167(4):323–60. Available from: https://linkinghub.elsevier.com/retrieve/pii/S0022519384710745

22. Holder BP, Liao LE, Simon P, Boivin G, Beauchemin CAA. Design considerations in building in silico equivalents of common experimental influenza virus assays. Autoimmunity [Internet]. 2011 Jun 19 [cited 2020 Aug 5];44(4):282–93. Available from: http://www.tandfonline.com/doi/full/10.3109/08916934.2011.523267

23. Saenz RA, Quinlivan M, Elton D, MacRae S, Blunden AS, Mumford JA, et al. Dynamics of Influenza Virus Infection and Pathology. J Virol [Internet]. 2010 Apr 15 [cited 2020 Dec 15];84(8):3974–83. Available from: http://jvi.asm.org/

24. Hancioglu B, Swigon D, Clermont G. A dynamical model of human immune response to influenza A virus infection. J Theor Biol. 2007 May 7;246(1):70–86.

25. Swat MH, Thomas GL, Belmonte JM, Shirinifard A, Hmeljak D, Glazier JA. Multi-Scale Modeling of Tissues Using CompuCell3D. In: Methods in Cell Biology. Academic Press Inc.; 2012. p. 325– 66.

26. Davis JW, Hardy JL. In Vitro Studies with Modoc Virus in Vero Cells: Plaque Assay and Kinetics of Growth, Neutralization, and Thermal Inactivation. Appl Environ Microbiol. 1973;26(3).

27. Kropinski AM, Mazzocco A, Waddell TE, Lingohr E, Johnson RP. Enumeration of bacteriophages by double agar overlay plaque assay. Methods Mol Biol [Internet]. 2009 [cited 2020 Nov 30];501:69–76. Available from: http://www.dsmz.de/}.

28. Tobita K. Permanent canine kidney (MDCK) cells for isolation and plaque assay of influenza B viruses. Med Microbiol Immunol [Internet]. 1975 Dec [cited 2020 Nov 30];162(1):23–7. Available from: https://link.springer.com/article/10.1007/BF02123574

29. Porterfield JS. A Simple Plaque Inhibition Test for Antiviral Agents: Application to Assay of Interferon. Lancet. 1959;326–7.

30. Rittenberg MB, Pratt KL. Antitrinitrophenyl (TNP) Plaque Assay. Primary Response of Balb/c Mice to Soluble and Particupre Immunogen. Exp Biol Med [Internet]. 1969 Nov 1 [cited 2020 Nov 30];132(2):575–81. Available from: http://ebm.sagepub.com/lookup/doi/10.3181/00379727-132-34264

31. Lindenmann J, Gifford GE. Studies on vaccinia virus plaque formation and its inhibition by interferon. III. A simplified plaque inhibition assay of interferon. Virology. 1963 Mar 1;19(3):302– 9.

32. Hayden FG, Cote KM, Douglas RG. Plaque inhibition assay for drug susceptibility testing of influenza viruses. Antimicrob Agents Chemother [Internet]. 1980 May 1 [cited 2020 Nov 30];17(5):865–70. Available from: http://aac.asm.org/

33. Weaver JJA, Shoemaker JE. Mathematical Modeling of RNA Virus Sensing Pathways Reveals Paracrine Signaling as the Primary Factor Regulating Excessive Cytokine Production. Processes [Internet]. 2020 Jun 20 [cited 2020 Aug 5];8(6):719. Available from: https://www.mdpi.com/2227-9717/8/6/719

34. Smith AM, Perelson AS. Influenza A virus infection kinetics: Quantitative data and models [Internet]. Vol. 3, Wiley Interdisciplinary Reviews: Systems Biology and Medicine. NIH Public Access; 2011 [cited 2020 Aug 5]. p. 429–45. Available from: /pmc/articles/PMC3256983/?report=abstract

35. Sun L, Liu S, Chen ZJ. SnapShot: Pathways of Antiviral Innate Immunity. Cell [Internet]. 2010 Feb 5 [cited 2019 Sep 9];140(3):436-436.e2. Available from: https://www.sciencedirect.com/science/article/pii/S0092867410000760?via%3Dihub

36. Dou D, Revol R, Östbye H, Wang H, Daniels R. Influenza A Virus Cell Entry, Replication, Virion Assembly and Movement. Front Immunol [Internet]. 2018 [cited 2019 Sep 9];9:1581. Available from: http://www.ncbi.nlm.nih.gov/pubmed/30079062

37. Frieman M, Heise M, Baric R. SARS coronavirus and innate immunity. Virus Res. 2008 Apr 1;133(1):101–12.

38. Opitz B, Rejaibi A, Dauber B, Eckhard J, Vinzing M, Schmeck B, et al. IFN? induction by influenza A virus is mediated by RIG-I which is regulated by the viral NS1 protein. Cell Microbiol [Internet]. 2007 Apr 1 [cited 2019 Sep 9];9(4):930–8. Available from: http://doi.wiley.com/10.1111/j.1462-5822.2006.00841.x

39. Wu W, Zhang W, Duggan ES, Booth JL, Zou MH, Metcalf JP. RIG-I and TLR3 are both required for maximum interferon induction by influenza virus in human lung alveolar epithelial cells. Virology. 2015 Aug 1;482:181–8.

40. Fujita T, Onoguchi K, Onomoto K, Hirai R, Yoneyama M. Triggering antiviral response by RIG-I-related RNA helicases. Biochimie [Internet]. 2007 Jun 1 [cited 2019 Sep 24];89(6–7):754–60. Available from: https://www.sciencedirect.com/science/article/abs/pii/S0300908407000247?via%3Dihub

41. Gack MU, Albrecht RA, Urano T, Inn K-S, Huang I-C, Carnero E, et al. Influenza A Virus NS1 Targets the Ubiquitin Ligase TRIM25 to Evade Recognition by the Host Viral RNA Sensor RIG-I. Cell Host Microbe [Internet]. 2009 May 21 [cited 2019 Sep 9];5(5):439–49. Available from: https://www.sciencedirect.com/science/article/pii/S1931312809001073

42. Rajsbaum R, Albrecht RA, Wang MK, Maharaj NP, Versteeg GA,Nistal-Villán E, et al. Species-Specific Inhibition of RIG-I Ubiquitination and IFN Induction by the Influenza A Virus NS1 Protein. Pekosz A, editor. PLoS Pathog [Internet]. 2012 Nov 29 [cited 2019 Sep 9];8(11):e1003059. Available from: http://dx.plos.org/10.1371/journal.ppat.1003059

43. Yuan S, Peng L, Park JJ, Hu Y, Devarkar SC, Dong MB, et al. Nonstructural protein 1 of SARS-CoV-2 is a potent pathogenicity factor 1 redirecting host protein synthesis machinery toward viral RNA. 2 3. bioRxiv [Internet]. 2020 Aug 10 [cited 2020 Sep 18];2020.08.09.243451. Available from: https://doi.org/10.1101/2020.08.09.243451

44. Le Page C, Génin P, Baines MG, Hiscott J. Interferon activation and innate immunity. Rev Immunogenet [Internet]. 2000 [cited 2019 Sep 23];2(3):374–86. Available from: http://www.ncbi.nlm.nih.gov/pubmed/11256746

45. Veer MJ de, Holko M, Frevel M, Walker E, Der S, Paranjape JM, et al. Functional classification of interferon‐stimulated genes identified using microarrays. J Leukoc Biol [Internet]. 2001 Jun 1 [cited 2019 Sep 23];69(6):912–20. Available from: https://jlb.onlinelibrary.wiley.com/doi/full/10.1189/jlb.69.6.912?sid=nlm%3Apubmed

46. Schneider WM, Chevillotte MD, Rice CM. Interferon-stimulated genes: a complex web of host defenses. Annu Rev Immunol [Internet]. 2014 [cited 2019 Jul 3];32:513–45. Available from: http://www.ncbi.nlm.nih.gov/pubmed/24555472

47. Shapira SD, Gat-Viks I, Shum BO V, Dricot A, de Grace MM, Wu L, et al. A physical and regulatory map of host-influenza interactions reveals pathways in H1N1 infection. Cell [Internet]. 2009 Dec 24 [cited 2019 Jun 26];139(7):1255–67. Available from: http://www.ncbi.nlm.nih.gov/pubmed/20064372

48. Coppey M, Berezhkovskii AM, Sealfon SC, Shvartsman SY. Time and length scales of autocrine signals in three dimensions. Biophys J. 2007 Sep 15;93(6):1917–22.

49. Sharova L V., Sharov AA, Nedorezov T, Piao Y, Shaik N, Ko MSH. Database for mRNA half-life of 19 977 genes obtained by DNA microarray analysis of pluripotent and differentiating mouse embryonic stem cells. DNA Res. 2009 Feb;16(1):45–58.

50. Prakash A, Levy DE. Regulation of IRF7 through cell type-specific protein stability. Biochem Biophys Res Commun. 2006 Mar 31;342(1):50–6.

51. Cohen LS, Studzinski GP. Correlation between cell enlargement and nucleic acid and protein content of hela cells in unbalanced growth produced by inhibitors of DNA synthesis. J Cell Physiol [Internet]. 1967 Jun 1 [cited 2020 Dec 7];69(3):331–9. Available from: http://doi.wiley.com/10.1002/jcp.1040690309

52. Cambridge SB, Gnad F, Nguyen C, Bermejo JL, Krüger M, Mann M. Systems-wide proteomic analysis in mammalian cells reveals conserved, functional protein turnover. J Proteome Res. 2011 Dec 2;10(12):5275–84.

53. Shapira SD, Gat-Viks I, Shum BOV, Dricot A, de Grace MM, Wu L, et al. A Physical and Regulatory Map of Host-Influenza Interactions Reveals Pathways in H1N1 Infection. Cell [Internet]. 2009 Dec 24 [cited 2020 Feb 10];139(7):1255–67. Available from: http://www.ncbi.nlm.nih.gov/pubmed/20064372

54. Ding Y, Chen L, Wu W, Yang J, Yang Z, Liu S. Andrographolide inhibits influenza A virus-induced inflammation in a murine model through NF-κB and JAK-STAT signaling pathway. Microbes Infect. 2017 Dec 1;19(12):605–15.

55. Lin R-J, Liao C-L, Lin E, Lin Y-L. Blocking of the Alpha Interferon-Induced Jak-Stat Signaling Pathway by Japanese Encephalitis Virus Infection. J Virol [Internet]. 2004 Sep 1 [cited 2021 Jan 7];78(17):9285–94. Available from: http://jvi.asm.org/

56. Demedts IK, Bracke KR, Maes T, Joos GF, Brusselle GG. Different roles for human lung dendritic cell subsets in pulmonary immune defense mechanisms. Am J Respir Cell Mol Biol [Internet]. 2006 Sep 20 [cited 2020 Nov 6];35(3):387–93. Available from: http://www.atsjournals.org/doi/abs/10.1165/rcmb.2005-0382OC

57. COVID-19 - Synairgen [Internet]. [cited 2020 Oct 2]. Available from: https://www.synairgen.com/covid-19/

58. Mantlo E, Bukreyeva N, Maruyama J, Paessler S, Huang C. Antiviral activities of type I interferons to SARS-CoV-2 infection. Antiviral Res [Internet]. 2020 Jul 1 [cited 2020 Oct 2];179:104811. Available from: /pmc/articles/PMC7188648/?report=abstract

59. Diebold SS, Kaisho T, Hemmi H, Akira S, Reis e Sousa C. Innate antiviral responses by means of TLR7-mediated recognition of single-stranded RNA. Science [Internet]. 2004 Mar 5 [cited 2019 Sep 24];303(5663):1529–31. Available from: http://www.ncbi.nlm.nih.gov/pubmed/14976261

